# Testing for positive selection in multi-copy gene families using a reconciliation approach

**DOI:** 10.1101/2025.08.18.670524

**Authors:** Yoann Pellen, Eyal Privman

**Affiliations:** Institute of Evolution, Department of Evolutionary and Environmental Biology, University of Haifa, Israel

## Abstract

Multi-copy gene families evolve through dynamic processes of duplication, loss, and sequence divergence, often exhibiting complex paralogy and orthology relationships that complicate the detection of adaptive evolution. Traditional dN/dS-based positive selection analyses are typically limited to single-copy genes, overlooking most of the adaptive signal in multi-copy families. We present a novel pipeline that integrates existing tools to map branch-specific positive selection events from gene trees onto species phylogenies. Our approach incorporates a normalization procedure that corrects for biases caused by variable branch lengths, reducing false enrichment signals on longer branches. Applied to the odorant receptor repertoires of ants, the framework successfully identified branches of the species tree enriched for positive selection, revealing clade-specific adaptive patterns obscured by conventional methods. This reconciliation-based strategy enables detection of adaptive hotspots in complex gene families, providing a versatile tool for linking molecular evolution to species-level diversification.

## Introduction

Understanding how genes evolve, and how their evolution shapes organismal traits, is a central goal of evolutionary biology. Gene evolution includes gene duplications, losses, and sequence divergence. Natural selection may act on duplicated genes as their sequences mutate, favouring adaptive genetic variants. Inferring adaptive evolution requires accurate phylogenetic reconstruction and comparative sequence analyses across species and gene families. A common approach to infer adaptive evolution on protein coding genes is the dN/dS (ω) ratio, which compares the rate of nonsynonymous (amino acid changing) to synonymous (silent) substitutions. Values of ω > 1 represent positive selection driving changes in the protein, which is interpreted as adaptive evolution. Elaborate models and tests were developed, such as the branch-site test for positive selection (Zhang 2005), which allow the detection of episodic selection affecting specific branches of a gene tree. These approaches depend on reliable alignments, robust phylogenies, and correct assumptions about evolutionary history.

A predominant model describing the evolution of multi-copy gene families is the birth-and-death model of gene evolution. First articulated by Ohno (1970), it poses that gene families expand through gene duplication events (“births”) and contract through pseudogenization and deletion (“deaths”). Over time, this dynamic process generates gene repertoires that are lineage-specific and functionally diversified. In contrast to single-copy gene families where one-to-one orthologs can be found across distantly related species, genes evolving under the birth-and-death model frequently exhibit complex paralogy (genes related by duplication) and orthology relationships across species. Zhang (2003) demonstrated that duplicated genes often experience relaxed selection initially, followed by either neofunctionalization (the evolution of a novel function), sub-functionalization (the division of multiple functions among paralogs), or loss. The birth-and-death process is now recognized as a key mechanism shaping the diversity of gene families involved in environmental sensing, immunity, and development (Demuth and Hahn 2009; Mendivil Ramos and Ferrier 2012).

Several large gene families expanded dramatically through duplications, like the cytochrome P450 superfamily. Gene duplication of cytochrome P450 genes that code for metabolic enzymes played a crucial role in increasing functional diversity, enabling organisms to adapt to changing environmental conditions by metabolizing a broader range of endogenous and exogenous compounds. These duplications allow for neofunctionalization or subfunctionalization, enhancing biochemical specialization and regulatory complexity. Such expansions have been linked to ecological adaptations in diverse animal species, including resistance to toxins and specialized diets (Nebert et al. 1989; Sezutsu et al. 2013; Yu et al. 2017; Kondo et al. 2022). Kinases are another example of a superfamily that expanded early in eukaryotic evolution to facilitate more complex cellular signalling pathways (Manning et al. 2002; Brunet et al. 2016).

Odorant receptors (ORs) are one of the largest gene superfamilies, with over 1000 functional genes in rodents (Niimura and Nei 2007). The diversity of animal ORs is key to their capacity to interact with their environment, including social interactions with conspecifics. Their vital and diverse functions drove their dynamic evolution across diverse vertebrate and arthropod, and even gastropod species (Niimura 2012; Engsontia et al. 2015; Branstetter et al. 2017; Eyun et al. 2017; Rondón Guerrero et al. 2024). Many studies have compared the OR repertoire between humans and mice, revealing functional similarities and evolutionary relationships between them, as well as the massive loss of more than half of the ORs in the human lineage (Lapidot et al. 2001; Mandairon et al. 2009). The numbers of ORs are also highly diverse among insect species. Most notably, ORs were dramatically expanded in ants, which was attributed to the evolution of sociality and complex chemical communication. The vast majority of the OR genes in different ant species are quite young, having arisen by gene duplications since the evolutionary split between ants and their closest relatives (Zhou et al. 2012; Engsontia et al. 2015; McKenzie et al. 2016; Branstetter et al. 2017; McKenzie and Kronauer 2018).

However, evolutionary relationships inferred from individual genes often conflict with the species phylogeny, making it difficult to link evolution of adaptive phenotypes of species to adaptive evolution of their gene as detected by the inference of positive selection. Gene tree–species tree reconciliation provides a framework to systematically map gene tree branches onto species tree branches by accounting for evolutionary events that can cause topological discordance, including duplications and losses (Karlsson and Stenlid 2008; Demuth and Hahn 2009; Freitas and Nery 2020; Legan et al. 2021; Marlétaz et al. 2023; Niimura et al. 2024; Page and Charleston 1998). Reconciliation has been used to inform the comparison of expression patterns of ancestral and novel forms of a gene (Marlétaz et al. 2023), infer the ancestral gene numbers of large gene families along the phylogeny and correlate the gains and losses with species evolution (Niimura et al. 2024), and even correlate these events with gene families which copy numbers are known to vary with adaptive evolution (Wapinski et al. 2007; Karlsson and Stenlid 2008).

In the literature, we see two main approaches to study the molecular evolutionary basis of species adaptation, either focusing on single or multi-copy gene families. Single-copy genes are easier to work with as each gene in the dataset represents a species and the gene tree is assumed to be the same as the species tree. This is the common approach of studies using dN/dS tests for positive selection, as it allows to report genome-wide frequency of positive selection directly to species tree branches (e.g. Kosiol et al. 2008; Hawkins et al. 2019; Romiguier et al. 2022). This approach is simple, but more susceptible to errors in tree reconstruction and incomplete lineage sorting. More importantly, this approach is limited to the most conserved gene families, and so it is likely to miss much of the adaptive evolution in genomes.

The other common approach applied to multi-copy genes is to infer gene family expansions along specific species tree branches, which are interpreted as adaptive evolution (Hahn et al. 2007; Han et al. 2009; Schönknecht et al. 2013). Alas, this approach is limited to the use of gene numbers and does not account for sequence evolution. Ideally, dN/dS tests for positive selection would be used to infer positive selection in various branches of the gene trees. But such an approach is prohibitively challenging due to the need to map gene tree branches to the species phylogeny, which is complicated by the difficulty to resolve orthologs and paralogs. Horizontal gene transfer also comes into play and adds to this challenge, especially in prokaryotic species for which this phenomena has a major impact on the gene repertoire (Koonin et al. 2001).

Reconciling such gene trees with their species tree is challenging. One of the tools developed for this purpose is *GeneRax* (Morel et al. 2020), primarily used to identify gene duplications, losses and transfers on a species phylogeny. Here we used it for a different purpose – to map positive selection events from the gene tree of large gene families to the species phylogeny. We developed a method allowing to study large gene families evolving by birth-death model, using existing tools. We illustrate the application of our method using OR genes, the largest insect gene family, with extensive evidence for positive selection.

## Methods

The first part of the pipeline is the standard protocol used to test for positive selection on every branch of the gene tree, using *Guidance2* to build and mask multiple sequence alignments, *RAxML* for gene tree reconstruction, and the branch-site test for positive selection implemented in *Godon*. Then the second part uses *GeneRax* for gene-tree species-tree reconciliation. The reconciliation allows testing for enrichment of positive selection in gene tree branches that map to a specific species tree branch of interest, which we want to test for adaptation in the evolution of some species.

### Multiple sequence alignment and gene tree reconstruction

*Guidance2* (Sela et al. 2015) was run with *PRANK* (Löytynoja et al. 2012) as the alignment algorithm. *Guidance2* detects unreliably aligned regions in a multiple sequence alignment and allows masking specific amino acids/codons, instead of the common practice of removing whole columns. Thereby we eliminate the unreliable portions of the data, without removing more than necessary. We start with a masking cutoff of 0.8, however, if more than 20% of the alignment was masked, we lowered the cutoff by 0.1 until reaching less than 20% masked residues. *RAxML* (Stamatakis 2014) was used to build gene trees based on the masked amino acid alignment using the PROTCATLG model.

### Positive selection inference

All gene tree branches were tested using the branch-site test for positive selection (Zhang 2005) implemented in *Godon* (Davydov et al. 2019), with branch lengths estimated by the M0 model, with 4 categories for codon rate variation. A likelihood ratio test was conducted for each branch contrasting between the null model, not allowing positive selection (dN/dS ≤ 1), and the alternative model, allowing for positive selection (additional site category with dN/dS > 1). The likelihood ratio test p-values were then corrected for multiple testing using the Benjamini-Hochberg (1995) method.

### Reconciliation-based enrichment test for positive selection

Reconciliation between the gene trees and the species tree was done using *GeneRax* (version 1.1.0, Morel et al., 2020), with the UndatedDL reconciliation model. This allows mapping gene tree branches that were tested for positive selection to the species tree branch that we wish to test for enrichment.

### Using GeneRax to map gene tree branches on the species tree

As input, *GeneRax* requires a rooted species tree in Newick format, a set of aligned gene family sequences in FASTA or PHYLIP format, and optionally, user-provided gene trees for each family. It produces a set of reconciled gene trees, each annotated with evolutionary events such as duplications, transfers, and losses. It is commonly used to detect and map such events to the phylogeny (Smith and Hahn 2021; Goulty et al. 2023; Marlétaz et al. 2023). The authors recommend visualizing *GeneRax* results using tools such as *ThirdKind* (Penel et al. 2022), a dedicated tool designed to graphically display reconciled trees. *ThirdKind* takes as input the “.*reconciled*.*xml”* files produced by *GeneRax* and renders dual representations of the gene tree and species tree, including graphical overlays of evolutionary events like gene duplications (marked as squares) and transfers (arrows).

While producing visual output is the most common use case of the *GeneRax*, for our purposes we extend its application by parsing the “.*reconciled*.*xml”* output to extract gene-to-species branch mapping. The “.*reconciled*.*xml”* file is in two parts: the first one is simply the species tree, listing all branches and leaves. It is the second part which is useful to us, with the mapping of gene tree branches on species tree branches (Figure 2). We extract associations between gene tree branches and species tree branches using individual labelling of every internal branch. Labelling of internal branches in the species tree can be done by running generax.py -f treelabel -it <input species tree file name> -ot <output species tree file name>. The internal branches labelling for the gene trees is automatically done when running *generax*.*sh*.

**Figure 1:**
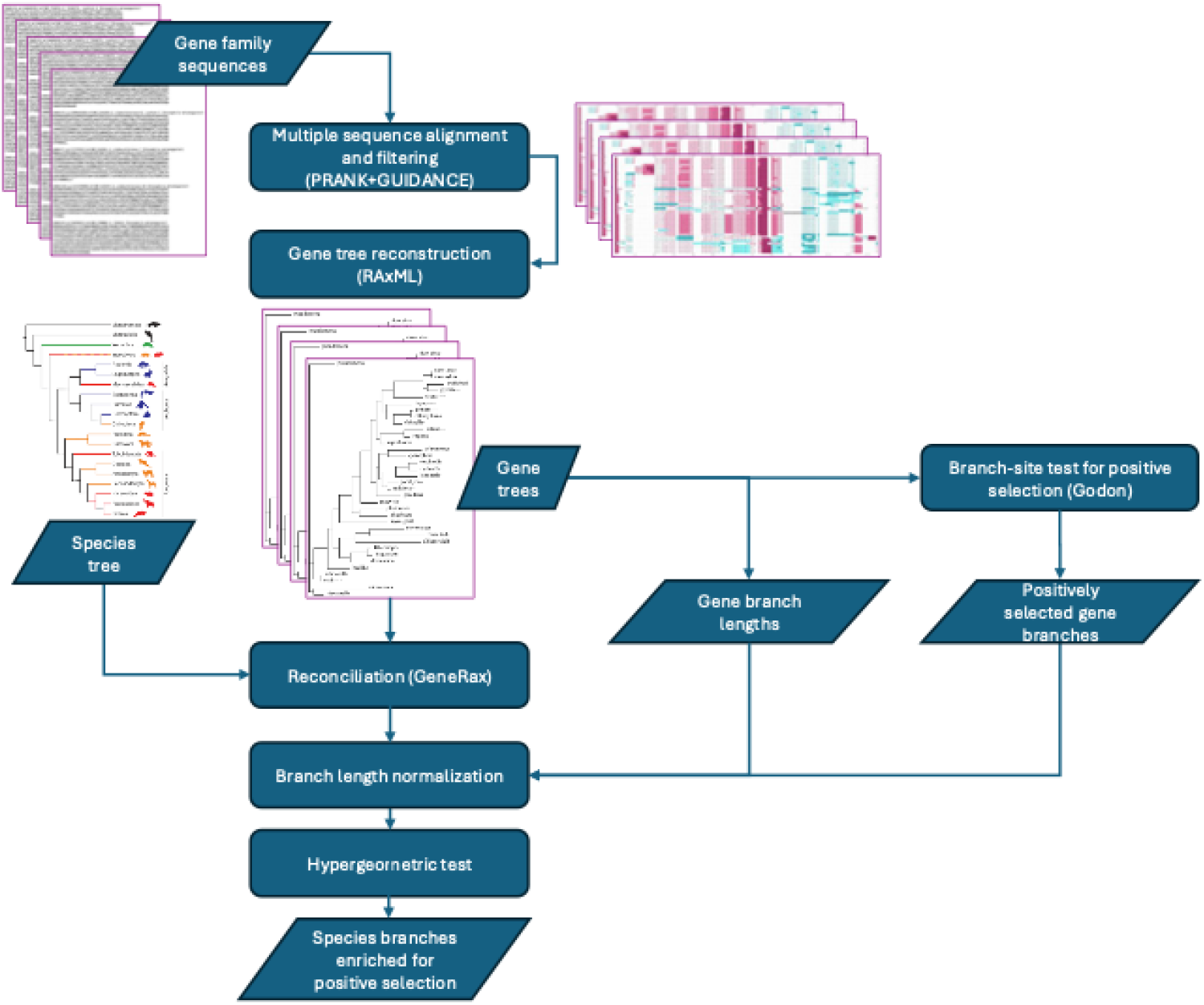
The positive selection reconciliation pipeline.

**Figure 2:**
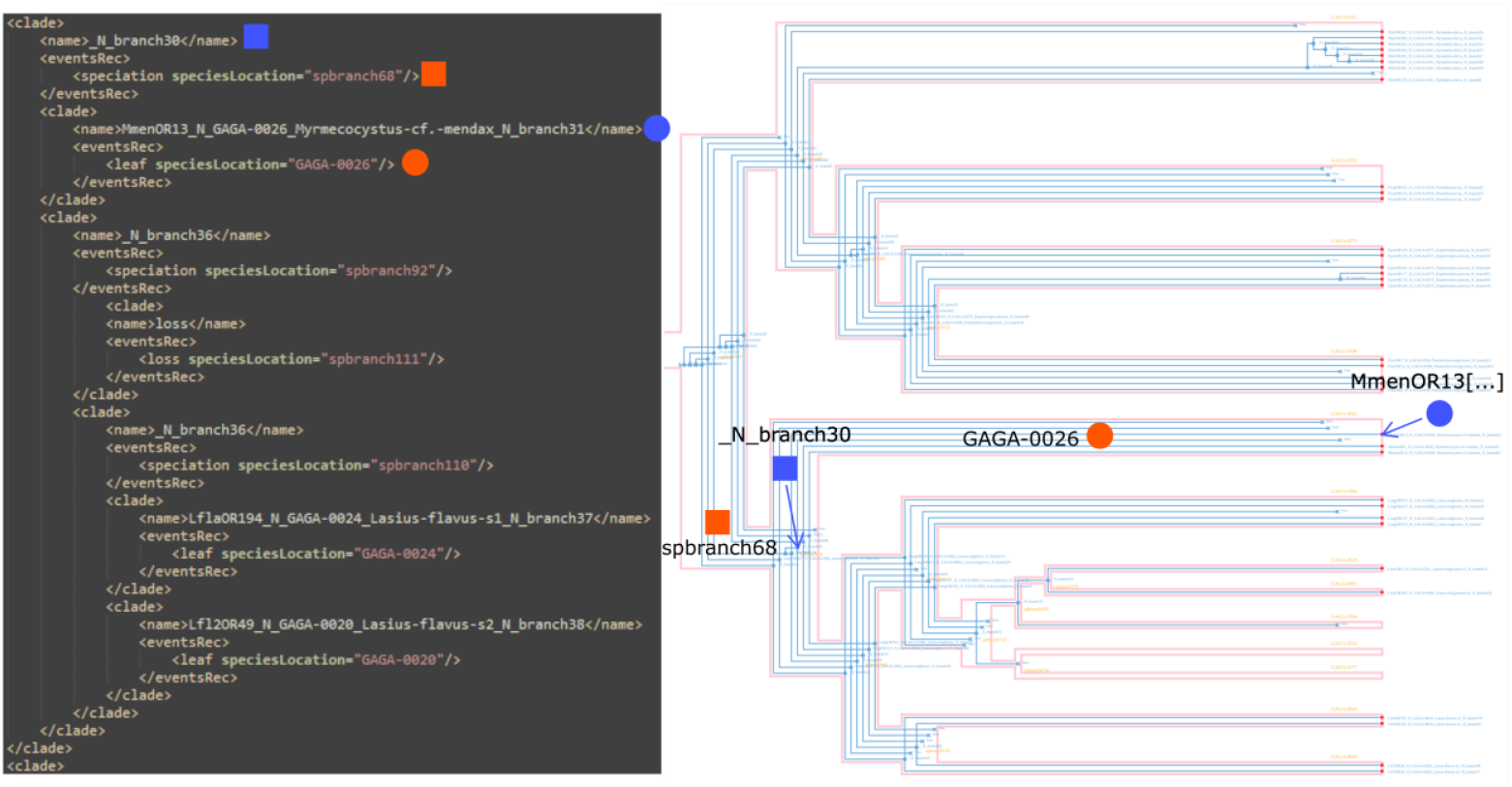
Part of the XML output of *GeneRax* (left), and its graphical visualization in *ThirdKind* (right). Gene tree branches (in blue) are marked with their corresponding species tree branch (in red) using the same shape.

From there, every branch and leaf from the gene tree is listed with the tag “*<name>*“, and its mapping on the species tree is specified with the tag “*speciesLocation=*“. Going through the file line by line, we match every gene tree branch to a species tree branch. For example, the gene branch “_N_branch30” (blue square) is mapped on the species branch “spbranch68” (red square). In another example, the gene branch “MmenOR13[…]” (blue circle), which is a leaf of the gene tree, is mapped on the species branch “GAGA-0026” (red circle), which is a leaf of the species tree. This mapping is done when running the script *generax*.*sh*.

### Hypergeometric test for enrichment of positive selection

We then used a hypergeometric test, implemented in the python module *SciPy* (Virtanen et al. 2020), to test for enrichment of positive selection in the species tree branches of interest. The hypergeometric distribution is a discrete probability distribution of *X* successes (random draws for which the object drawn has some desired feature, in this case positive selection) in *N* draws without replacement from a finite population of size *M* that contains exactly *n* objects with that feature (i.e. successes). Taking p-values under 5% from *Godon* as our successes, our population size *M* is the total number of gene branches in our dataset, and the number of draws *N* is the number of gene branches mapped on the species branch of interest.

Importantly, we addressed a critical bias in tests of this kind: longer branches have more opportunities to accumulate substitutions, and thus a greater chance of showing evidence for positive selection. Without correction, this would lead to an overrepresentation of positive selection signals on longer branches, which could be mistaken as biologically meaningful signals associated with the evolution of novel traits when they may simply reflect longer evolutionary time.

To correct for this issue, we decided to standardise the population parameter instead of simply taking the count of the number of branches. This normalization ensures that species-tree branches with longer evolutionary histories do not appear to be enriched for positive selection merely due to longer evolutionary time.

### Correcting for branch length

Our first approach was to try to sum the branches length instead of simply counting them for the hypergeometric test as presented above. We summed the length of all the gene branches for both the population (*M*) and sample (*N*) sizes. In a third approach, we normalized the number of branches by a factor defined by their branch length relative to the average gene tree branch length of the studied clade. To do so, for each branch of interest of the species tree, we normalized the length of every gene branch by dividing it by the average gene branch length in the clade, before summing all of them. All scripts are available on GitHub (https://github.com/yoannpellen/GAGA_ORs), with a detailed description on how to run them.

### Application to real data

This method was applied in Pellen et al. (2025) to a dataset of 50,657 OR sequences that were annotated in genomes of 145 ant species (Vizueta et al. 2025). Due to the limits of the bioinformatic tools in our pipeline in dealing with such large numbers of sequences, we restricted the analysis to smaller clades of a dozen species at most, including transitions in our traits of interest. For the same reason, the OR superfamily was also divided into 30 subfamilies, and the results from all subfamilies were then pooled back together for each clade for the hypergeometric test. Here we show the results for a subset of branches, selected to represent the variability we saw in our results.

## Results and discussion

We observed high variability in the results depending on the method used (Table 1, Figure 3), both in terms of p-value but also in the observed fold-change in the proportion of branches with positive selection (enrichment or depletion). Without normalizing, we assessed enrichment by counting the number of gene tree branches mapped to a species tree branch that were found to be under positive selection. This method assumes all branches have equal opportunity to accumulate substitutions and hence equal chances of showing positive selection. The hypergeometric test gave statistically significant support (p-value < 0.05) for depletion of positives selection in branches 2 and 3, and a marginally significant depletion in branch 1 (p-value = 0.083).

**Table 1:**
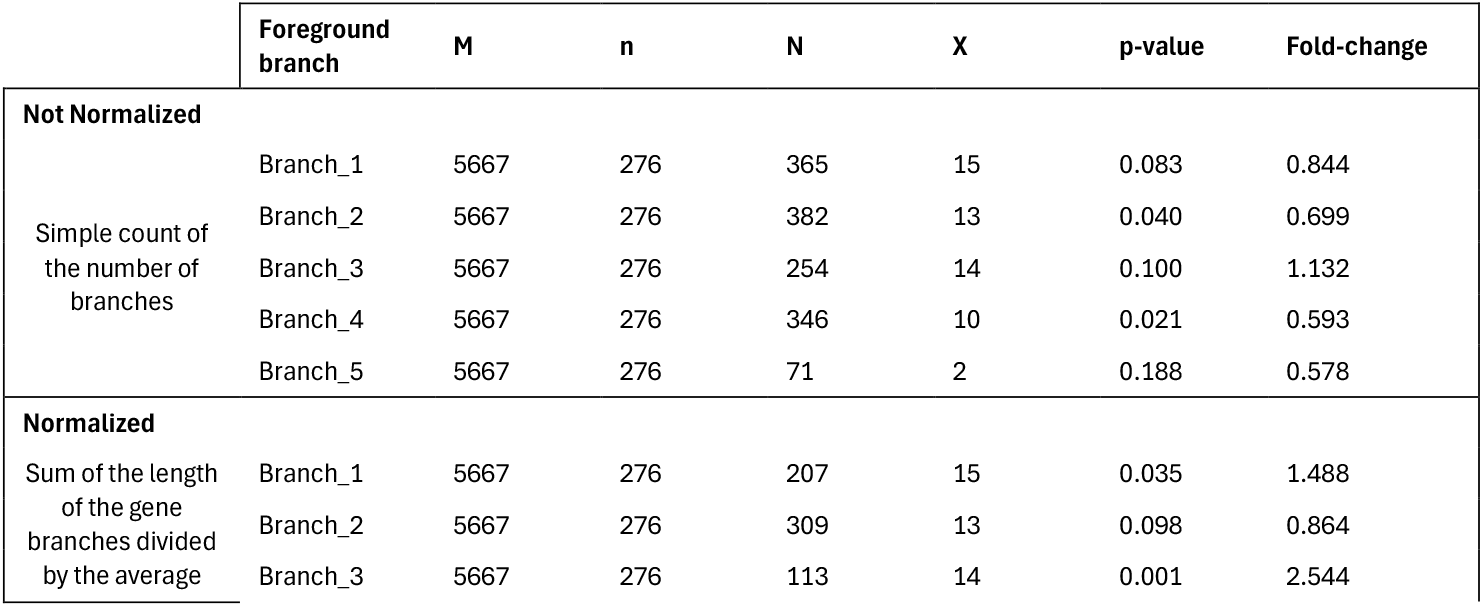
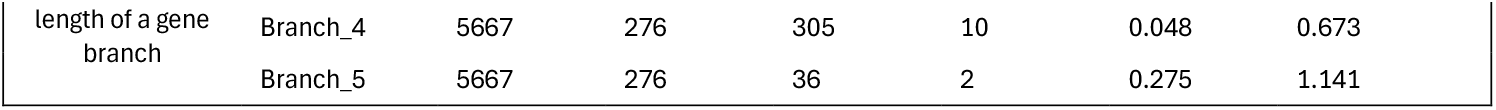
Count data used in the normalized and un-normalized versions of the tests for enrichment of positive selection.

**Figure 3:**
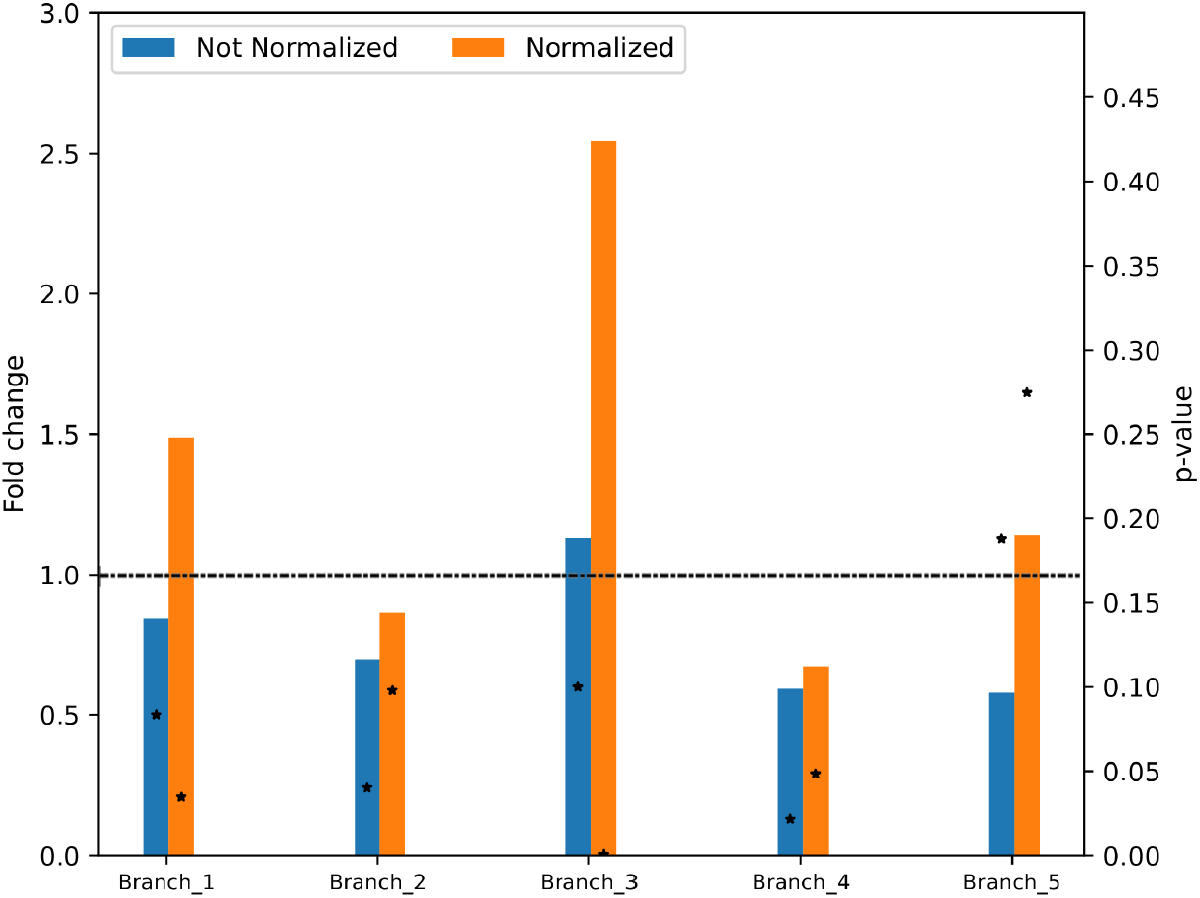
Comparison of the normalized vs. un-normalized versions of the tests for enrichment of positive selection. The plot shows differences in fold-change and p-value of the hypergeometric test. The dashed line marks a fold-change of one; values above one represent enrichment and values below one represent depletion of positive selection. Respective p-values are reported with black stars.

The un-normalized method assumes equal substitution opportunity across all branches, which may be biologically unrealistic and lead to a bias favoring longer branches. To address this bias, one may use units of branch lengths instead of counts of branches. We could use the sum of the gene tree branch lengths rather than their count. However, this length-based adjustment method is not applicable this context. It would results in small sums due to the short gene branch lengths, which are typically much shorter than 1 unit of evolutionary distance (the average gene branch length in this dataset was 0.063). This reduces the population size (*N*) to the point where it is smaller than the number of successes (*X*).

It is sensible to normalize the branches relative to the average branch length of the gene tree. This version of the enrichment test gave substantially different results. For instance, branch 1 became more significant (p-value = 0.035), but the direction of the fold-change was also impacted, showing an enrichment instead of depletion (fold-change = 1.488). The results for branch 2, despite still showing a depletion, lost their significance (p-value = 0.098). Branch 3 showed a much stronger enrichment (fold-change = 2.544, p-value = 0.001). Branches 4 and 5 showed little change, with branch 4 still harboring a significant enrichment and no significant result for branch 5.

The choice of normalization method substantially impacts both the detection and the direction of inferred enrichment or depletion in positive selection. Simply counting the number of branches inferred to be under positive selection is prone to false positives in longer branches. The length-based adjustment is inapplicable due to short gene tree branch lengths. The normalization method provides a solution, reflecting relative evolutionary opportunity, thereby avoiding statistical artifacts. It normalizes the number of gene tree branches by a factor defined as their branch length relative to the average gene tree branch length of the studied clade. This approach uses the average branch length as a unit of evolutionary time, which corresponds to a certain expectation regarding the probability for positive selection.

Understanding adaptive evolution within gene families shaped by complex evolutionary processes, including duplication, loss, and sequence divergence, requires methodological advances that go beyond the common single-copy gene family studies. In this study, we developed a pipeline that integrates existing tools in a novel approach for the study of multi-copy gene families, which can reveal the enrichment of positive selection in species tree branches. By combining tested tools (*Guidance2, RAxML, Godon*, and *GeneRax*), we established a robust framework to map gene tree-level selection signals onto species-level branches.

A key innovation of our approach lies in the reconciliation-based mapping of selection signals, which addresses the common issue of gene tree-species tree discordance in multi-copy gene families. The challenge of topological mismatch, due to either incomplete lineage sorting, gene duplication and loss, or horizontal gene transfer, limits the utility of standard dN/dS analysis pipelines to cases with congruence between gene and species trees. Our reconciliation approach allows us to account for this discordance and obtain a more comprehensive understand of where adaptive events are concentrated in the evolutionary history of a species group, taking into consideration all gene families in the genome. Importantly, large gene families characterized by birth-and-death evolutionary processes are more dynamic and less conserved than single-copy gene families. Therefore, such gene families are likely to be more important for adaptive evolution of species.

## Conclusion

This study presents a framework for detecting adaptive evolution in multi-copy gene families, using reconciliation-informed mapping of gene-level selection events to species-level branches. By using already existing tools in a novel way, we offer a solution to the longstanding problem of gene-tree species-tree discordance in molecular evolution analyses.

The methodology bridges a critical gap between molecular evolutionary inference and species-level trait evolution, offering a path toward more accurate identification of evolutionary hotspots of species adaptation. Importantly, it enables researchers to link gene family evolution, duplication patterns, and adaptive signatures across the tree of life, with potential applications in fields such as evolution of immune systems, metabolic adaptation, and sensory diversification.

With increasing availability of high-quality genomes and the ongoing refinement of phylogenomic tools, our approach provides a foundation for future evolutionary genomic studies aiming to contextualize molecular adaptation within the species phylogeny.

